# Collective decision making by rational agents with differing preferences

**DOI:** 10.1101/2020.01.13.904490

**Authors:** Richard P. Mann

## Abstract

Collective decisions can emerge from individual-level interactions between members of a group. These interactions are often seen as social feedback rules, whereby individuals copy the decisions they observe others making, creating a coherent group decision. The benefit of these behavioural rules to the individual agent can be understood as a transfer of information, whereby a focal individual learns about the world by gaining access to the information possessed by others. Previous studies have analysed this exchange of information by assuming that all agents share common goals. While differences in information and differences in preferences have often been conflated, little is known about how differences between agents’ underlying preferences affects the use and efficacy of social information. In this paper I develop a model of social information use by rational agents with differing preferences, and demonstrate that the resulting collective behaviour is strongly dependent on the structure of preference sharing within the group, as well as the quality of information in the environment. In particular, I show that strong social responses are expected by individuals that are habituated to noisy, uncertain environments where private information about the world is relatively weak. Furthermore, by investigating heterogeneous group structures I demonstrate a potential influence of cryptic minority subgroups that may illuminate the empirical link between personality and leadership.

## INTRODUCTION

The choices made by others serve as a conduit for information about the world that other individuals possess, and therefore enable a focal individual to make decisions with greater expected utility or fitness than they could do alone. This transmission of information, frequently labelled as ‘social information’, has been posited as an important factor driving collective cohesion in animal and human groups [1–3], alongside other benefits of aggregation such as dilution of predation risk [4].

The tendency of individuals to follow the decisions of others has been comprehensively demonstrated across many taxa (e.g. insects [5], fish [6, 7], birds [8, 9], and mammals [10–12], including humans [13, 14]). Many studies have posited simple social feedback rules as models for collective decision making, demonstrating how individual behavioural heuristics could generate cohesive collective decisions [15–17]. Other studies have sought to reveal the form of these social interaction rules via a data-driven approach [5, 18]. However, neither of these methods identify the source of the behavioural rules in terms of the direct benefit to the individual, an important evolutionary principle in unrelated groups [19]. More recent work has investigated how social interaction rules could be justified from the rational self-interest of the decisionmaking agents, either through an evolutionary analysis [20], or by direct calculation [1–3]. Identifying the individual fitness or utility motivations of social interactions is important for understanding how behaviour will vary with context, and thus to extrapolate from empirical observations to make predictions about social behaviour in other environments [3].

Studies that have sought to directly evaluate rational rules of social interaction [1–3] have addressed the fundamental question of social information: what can one individual learn by observing the choices made by others? The general approach to this question has centred on two conceptual points: (i) the choices of other agents are of interest because those agents have information that the focal individual lacks; and (ii) the focal agent can only infer what that information might be by considering why the other agent made the choice it did. In considering why another agent may have made the choice they did, the focal agent must have a model for how that agent responds to different information. A simple model is to assume that the other agent is identical to oneself, and therefore responds identically to any set of information [3]. However, groups are not, in general, composed of identical individuals, and agents may have different preferences based on their particular needs or tastes. This may be the case within same-species groups, as individuals can differ in, for example, age, genetic profile [21], or current nutritional needs. Even more clearly, groups may be composed of multiple species [8, 22], with potentially very different needs and goals between individuals from different species.

How then might agents make use of social information when their preferences differ? It is important here to distinguish between differences in preference and differences in information. Information and preference differences are frequently conflated in models inspired by physical systems, where both are typically expressed in terms of attractive forces or potentials (e.g. [23–26]), but in reality these are different sources of possible conflict, with quite different effects upon a rational agent. Consider two individuals, one of which is trained to associate food with red markings, the other with blue marking. This may result in an difference in revealed colour preference between the when both markings are displayed. However, this is colour preference is, from a rational perspective, a cipher for both individuals’ common preference for food. It is therefore better understood as a difference of information: the two individuals may differ in where they believe food to be located, but both still share a desire to find the food, and should therefore be interested in what the other knows. Conversely, two individuals with exactly the same information may make different choices if they have different goals, and be entirely uninterested in the choices each other make - for example a vegetarian and a committed carnivore choosing where to eat based on the same set of restaurant reviews. I have previously considered differences in information when agents have identical preferences [3], where agents can view each other as more or less direct proxies in terms of response to information. Differences in preferences matter because they change what one individual can learn from observing another, as the actions of those other agents may be different to what the focal agent would have done with the same information.

Here I develop a model for collective decision making by rational agents with differing preferences, which seek to maximise differing utility functions [27]. This is based on a model of utility structured as an array of environmental factors characterising different choices, which can encode differing degrees of correlation between individual preferences. Using this framework of utility functions I then derive the rational decision making rules for individuals to follow, assuming that individuals have common acquired knowledge about the expected range of environmental characteristics and the typical degree to which their preferences align. Finally I derive the expected behaviour of the individuals and the group when faced with a binary decision between two alternatives, A and B, which can represent different foraging patches, different movement directions or any other mutually exclusive activities. Following ref. [3], I evaluate the expected behaviour from the perspective of an external observer (as opposed to the agents themselves), and thus make predictions about the likely characteristics of collective decision making seen in empirical studies and in real human and animal groups.

## RESULTS

### Differing preferences induce environmental dependence in decision making

I evaluated the probability that a focal individual will choose option A, under natural conditions where the experimental noise level (*η*) matches the habitual noise level (*ϵ*), conditioned on different sequences of previous decisions, and within a group where individual preferences have a characteristic correlation *ρ*. I further calculated how these probabilities varied with the habitual environmental noise level *ϵ*. The results of these calculations are shown in Figure 1, with each line corresponding to a different sequence of up to three previous decisions. Each line is labelled by the sequence of previous decisions in chronological order; for example AAB represents the case where the most recent observed choice was B, preceded by two agents choosing A. The two panels corresponds to a different degree of preference alignment (*ρ*) between individuals. These results demonstrate a strong dependence of social influence on the habitual environmental noise level. For relatively low preference alignment (*ρ* = 0.5, panel A), increasing the noise level e has the effect of increasing the tendency for the focal individual to follow the majority of previous decision makers. When the environmental noise level is low, social responses are very weak and individuals tend to follow their own private information. In the second case where preferences are highly aligned (*ρ* = 0.9, panel B) the response to social information is strengthened across the range of *ϵ*, but an interesting secondary effect is seen in addition to this general trend: as the noise level *ϵ* is increased there is a transition between weakly following the majority opinion to strongly following the most recent decisions in cases where the most recent decision conflicts with the majority. This can be seen clearly in the crossing of the lines representing the decision sequences AAB and BBA in panel B. These results should be compared with those of ref. [3], which correspond to the special case of *ρ* = 1. In that case there is no dependence on environmental noise levels, and the most recent decision is always favoured.

**FIG. 1.**
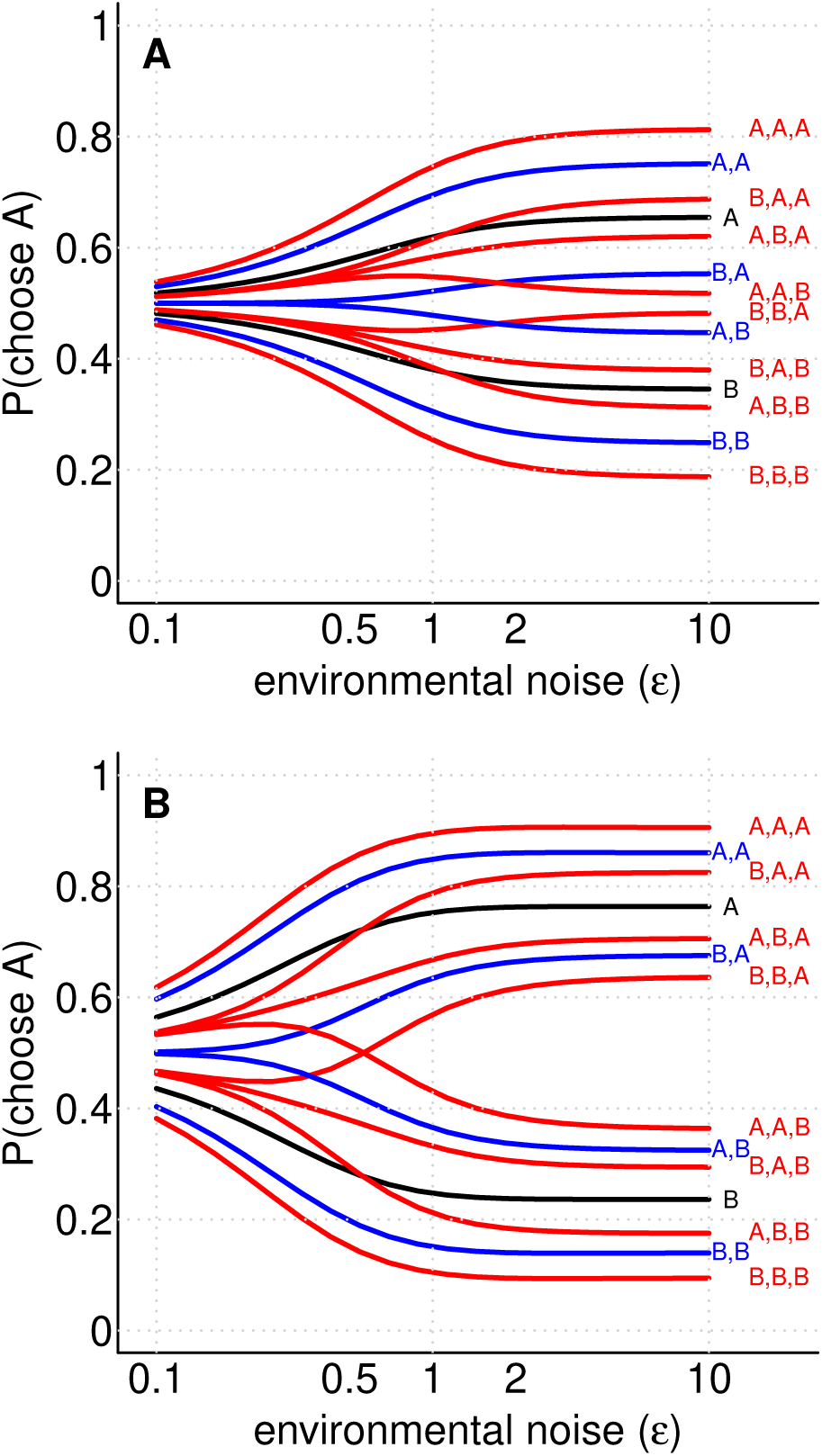
The probability for a focal agent to choose option A, conditioned on possible sequences of previously observed decisions by up to three other individuals, across a range of environmental noise levels, with weakly aligned preferences (*ρ* = 0.5, panel A) or strongly aligned preferences (*ρ* = 0.9, panel B).

### Relative social weighting

The theory described in the methods section results in a complex, recursive decision-making rule. However, a simpler perspective on the importance of the model parameters can be obtained if one focuses only on the decision faced by the second individual, who must balance the social information provided by the first decision against their own private information. How are these sources of information weighted? One can see from equation 12 that the private information of agent 2 enters in the second term, weighted by a factor of 1/*ϵ*, while the social information is contained in the third term and weighted by 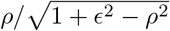. From the ratio of these two weightings we can define the ‘relative social weighting’ (RSW),

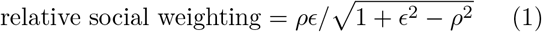

This relationship contains several important special cases. When individuals share identical preferences (ρ = 1), the choices made by others are a direct proxy for the decision the focal agent would have made themselves, and thus the RSW is one. When preferences are not identical (*ρ* < 1), private information dominates in cases where that information is reliable (*ϵ* << 1), and thus the RSW is close to zero. However, when private information is not reliable (*ϵ* >> 1), the choices of others are weighted in proportion to how strongly preferences are aligned (RSW ≃ *ρ*). If there is no correlation between two individual’s utility functions (*ρ* = 0) then the choices made by one convey no social information to the other, and the RSW is zero.

### Sequence ordering in previous decisions

I investigated further how preference alignment and environmental noise determine how a focal individual uses social information, focusing on the possible tension between a majority of previous decisions and the most recent of those decisions. To explore this tension I consider a simple conflict: a sequence of previous decisions of the form: BBA. That is, the focal individual is presented with two previous decisions in favour of B, followed by one in favour of A, such that the majority of previous decisions favour B, but the most recent social information favours A. For a range of values of *ϵ* and *ρ*, and for two different experimental conditions n, I calculated the probability that the focal individual will choose option A. The results, shown in Figure 2, indicate a consistent pattern for resolving this conflict. When tested under naturalistic conditions (*η*/*ϵ* = 1, panel A) individuals that are habituated to a noisy environment and have a high degree of preference alignment are more likely to follow the most recent decision maker. Conversely, individuals that are habituated to low noise environments and/or have a low degree of preference alignment, are more likely to follow the majority. The white contour line indicates where either outcome is equally likely, i.e. the agent follows the most recent decision 50% of the time. This effect is amplified by low-noise experimental conditions (e.g., *η*/*ϵ* = 1/2, panel B), but the transition contour of *P*(*A*) = 0.5 remains consistent across the different experimental treatments. In panel C the probability to choose A is plotted against the relative social weighting defined above, showing that this combination of *ρ* and *ϵ* explains much of the variation in the decision probability, with the most recent decision being favoured when the relative social weighting is above approximately 0.5.

**FIG. 2.**
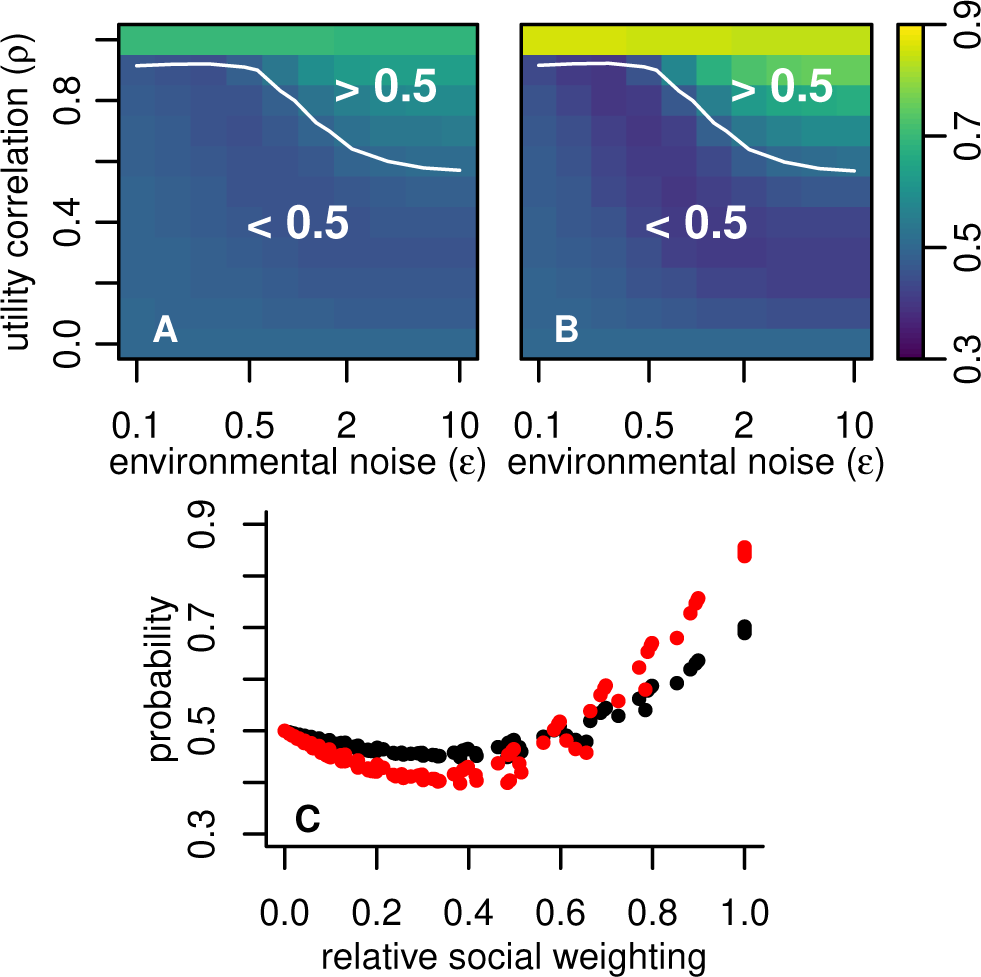
Resolving conflicts between majority decisions and recent decisions. Panel A shows the probability that a focal individual will select option A, conditioned on observing three previous decisions of the form: BBA, as a function of the utility function correlation and the environmental noise level, tested under natural conditions *η*/*ϵ* = 1). Panel B shows the equivalent probabilities, assuming the focal individual is tested under low-noise laboratory conditions (*η*/*ϵ* = 1/2). In both cases the focal agent favours recent information when utility alignment is strong and environmental noise is high. Panel C shows the results for the natural setting (black points) and the laboratory setting (red points) plotted against the relative social weighting

### Relative social weighting drives consensus

Having determined that preference alignment and environmental noise have a strong effect on how a focal individual makes use of social information, we need to understand the consequences of this for collective behaviour. Of particular interest is the degree to which groups can come to consensus decisions and remain cohesive.

Assuming a group of six individuals making sequential choices according to the rational decision-making rule, I evaluated the probability for each possible collective outcome in the binary decision-making scenario, in terms of the eventual number of individuals selecting option A, *n_A_* and those choosing option B, *n_B_*. This calculation considered the probability of every sequence of decisions that could give rise to a given collective outcome and summed over these to give the final probabilities. Each of these collective outcomes is given a ‘consensus score’, defined as:

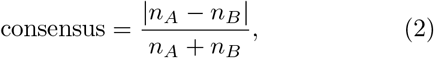

such that a consensus score of one indicates that all individuals chose the same option, and a consensus score of zero indicates an equal split between A and B.

I performed this calculation for the same range of environmental noise levels, preference alignments and experimental conditions as in the analysis of sequence ordering above. The results, shown in Figure 3, demonstrate that consensus depends strongly on these factors. When individuals are habituated to a noisy environment or have strongly aligned preferences then the group tends to come to a consensus decision, with a high consensus score under these conditions and in the naturalistic experimental environment (*η/ϵ* = 1, panel A). Conversely, consensus scores are low when the habitual environment has little noise, or when preferences have very low alignment. Low-noise experimental conditions (*η/ϵ* = 1/2) create greater consensus for any given values of *ρ* and *ϵ*, (panel B); this effect of experimental noise is in line with those explored in ref. [3]. In panel C the expected consensus score for each experimental condition is plotted as a function of the relative social weighting (black points for the *η/ϵ* = 1, red points for *η/ϵ* = 1/2). Here one can see that relative social weighting explains almost all the variation in consensus as a function of *ρ* and *ϵ*.

**FIG. 3.**
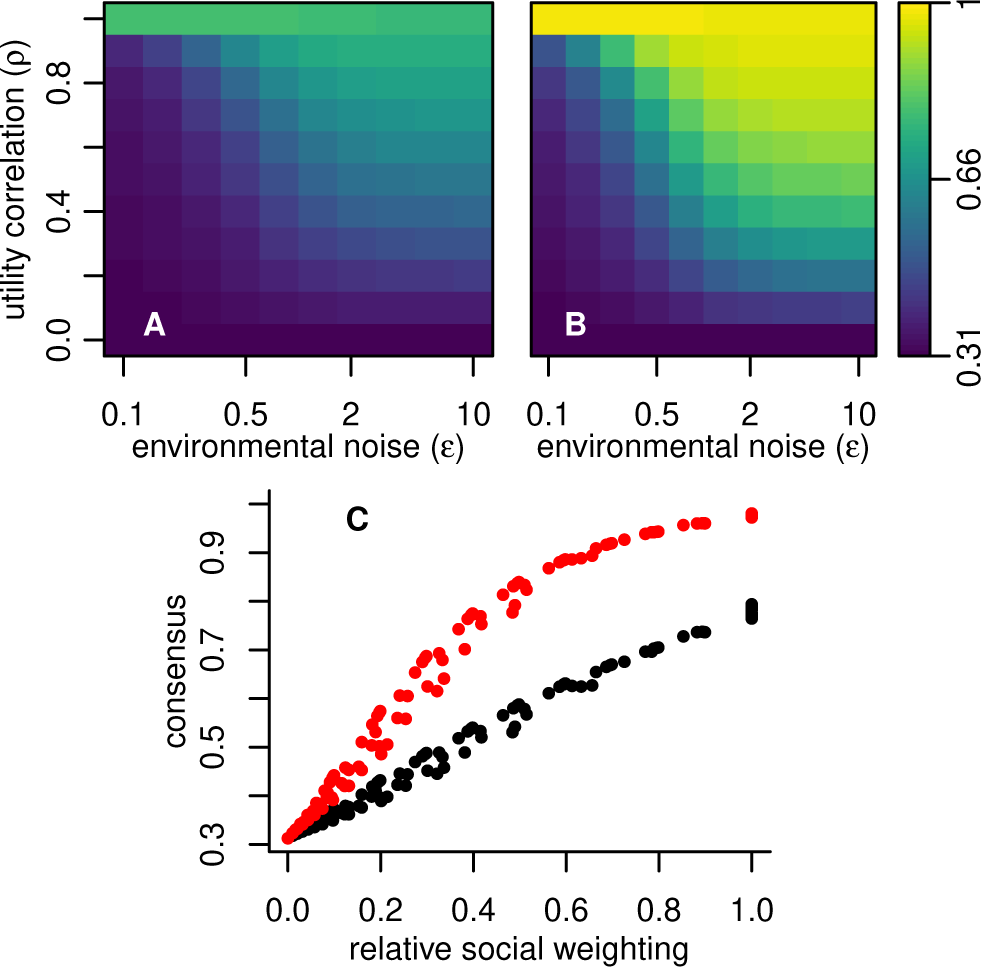
Expected group consensus as a function of environmental noise and utility correlation for six agents tested under naturalistic conditions (*η/ϵ* = 1, panel A) and low-noise laboratory conditions (*η/ϵ* = 1/2, panel B). Panel C shows the expected consensus for naturalistic (black points) and laboratory (red points) conditions as a function of the relative social weighting. Consensus is strong when relative social weighting is high and is enhanced by low-noise experimental conditions.

### The effect of cryptic group substructures

The analysis so far has looked at the behaviour of groups from a population that is undifferentiated, having a fixed utility correlation between any pair of individuals. Now I consider a group drawn from a population with further substructure: the existence of two distinct types of individual, *α* and *β*. I assume that individuals of the same type share a high correlation between their utility functions, whereas individuals of differing types have a low correlation. Furthermore, I assume that these individual types are cryptic, i.e. they are not directly observable by other individuals, except through decision-making behaviour. To illustrate the relevance of such a such a structure, consider a population with two distinct genotypes, with the result that individuals with each genotype require different relative amounts of carbohydrate and protein, but where the genotype has no other, directly-perceivable phenotypic effect. In such a population, we would expect the interactions between individuals to reflect the reality that some conspecifics more closely share their nutritional preferences than others.

First I consider a population of two equally-numerous types, with those of a given type having identical preferences (*ρ* = 1), and individuals of differing types having no correlation in their utility function (*ρ* = 0). I calculated the probability that a focal decision maker of type *α* will choose option A conditional on a variety of possible sequences of previous decisions, and for a range of possible environmental noise levels (by symmetry the results are the same for a focal individual of either type). These probabilities are shown in Figure 4A. These results show a superposition of two different responses to environmental noise. In the cases where all previous decision makers have made the same choice the probability for the focal individual to choose A changes little with environmental noise. This pattern is close to that seen for agents with identical preferences (see ref. [3] fig. 1). However, in cases where previous decisions have not been unanimous there is a strong dependence on the environmental noise, resembling that seen Figure 1A. The intuition behind this superposition is that when a focal agent observes all previous decision makers in agreement, they conclude that it is likely they are all of the same type, and thus share identical preferences. Conversely, observing disagreement implies a high probability that the previous decision makers were of differing types. The consequences of this are particularly striking for the case of low environmental noise: in such environments social influence is relatively low, except in the case of unanimity. That is, the difference between a unanimous set of previous decisions compared to a majority is qualitative rather than merely quantitative.

**FIG. 4.**
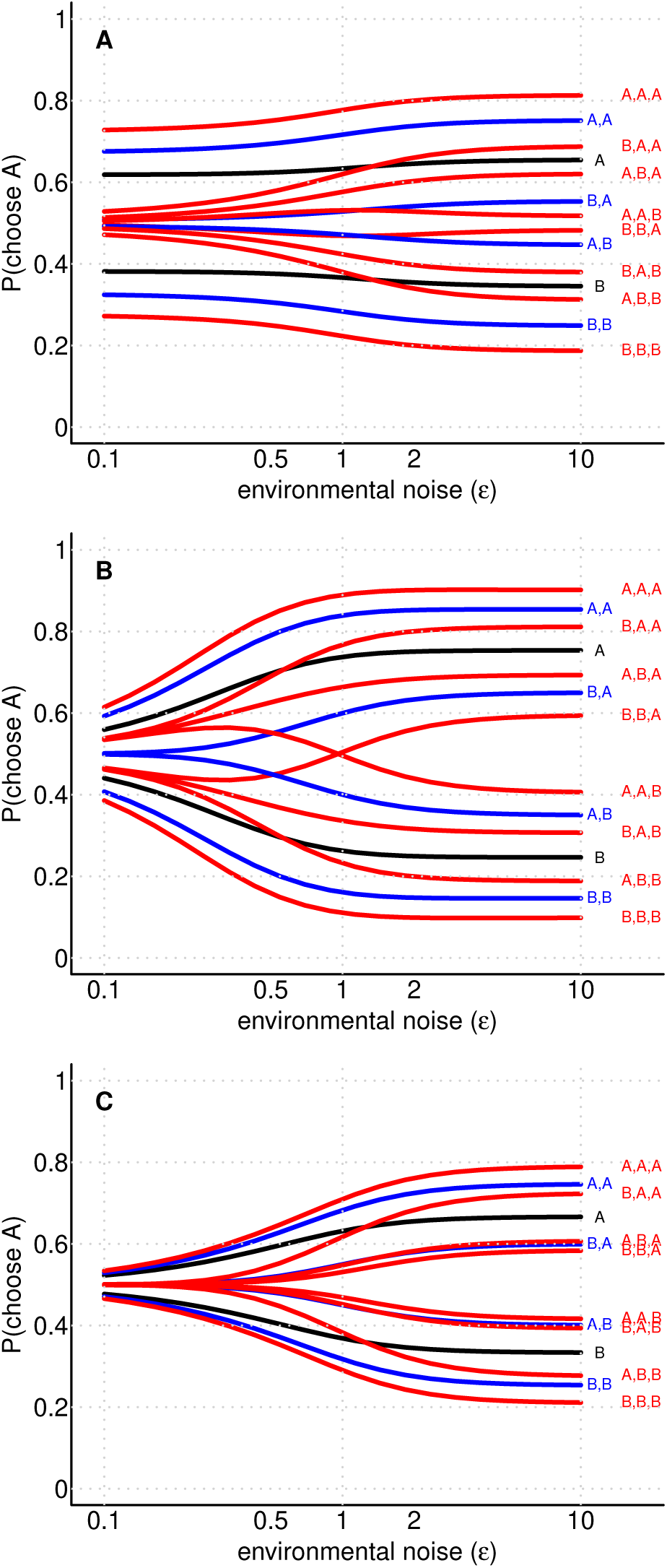
Scenarios with cryptic group substructures. Panel A shows the probability for a focal individual to choose option A as in Figure 1, when individuals are drawn from a population of two equally prevalent sub-types with identical within-type preferences and zero between-type correlation. Panels B and C show the equivalent probabilities for individuals of the majority type (panel B) and minority type (panel C) in the case where 90% of individuals are of the majority type, with high within-type utility alignment (*ρ* = 0.9) and low between-type correlation (*ρ* = 0.1)

In the example above the two distinct types were equally prevalent in the population as a whole. In general this will not be the case: frequency-dependent evolution can select for substantially unequal proportions of differing types of individual (e.g. [20]). What is the effect of such an imbalance on social behaviour? I considered a population composed of 90% type *α* individuals and 10% type *β*, where *ρ_α,α_* = *ρ_β,β_* = 0.9 and *ρ_α,β_* = 0.1. I calculated the probability for a focal agent of either type to choose option A, based on the same set of putative previous decision sequences as in the previous example. The results for an individual of the majority type *α* are shown in Fig. 4B, and those for an individual of the minority type *β* in Fig. 4C. From these plots one can see a substantial difference in the social behaviour of the two types of individual: type *α* individuals display strong social responses close to those of individuals in an undifferentiated and strongly aligned population (see Fig 1B), while type *β* individuals have a weak social response similar to those from an undifferentiated and weakly aligned population (see Fig 1A). This difference arises from the different utility correlation between individuals of each type and the population as a whole: those in the majority can assume that most other individuals have closely aligned preferences and therefore are worth following, while those in the minority understand that most other individuals have differing preferences and therefore convey little useful social information.

## DISCUSSION

In this paper I have described a model of collective decision making by rational individuals with differing preferences, utility functions or fitness outcomes when faced with decisions that depend on many factors. Developing this model has focused attention on precisely how the choices of one individual convey information about the likely utility of possible options to another individual, and when that information is likely to be reliable. This model goes beyond previous efforts to understand the foundations of social information by removing any assumption that individuals are identical, while distinguishing between differences in knowledge and differences of preference.

I have shown that the rational use of social information depends strongly upon the degree to which the focal decision maker believes that others share its preferences, with stronger social interactions between similar agents. This intuitive result provides an informational basis several for observed tendencies in animal groups, e.g. for the observed stronger social response to conspecifics compared to heterospecifics in mixed species bird groups [8, 22], for the tendency of baboons to follow movements initiated by close social affiliates [28] and for the preference of true conspecifics over a robotic imitation in zebrafish [29]. It is also likely to contribute to homophily in human societies, for example in housing choice [30], where the presence of many households with similar characteristics to oneself in an area provides useful information that your own needs and preferences can be met locally. In this way collective patterns such as neighbourhood segregation may be driven by the different information values of other individuals as well as other explanations such as the dislike of being in a local minority [31].

The use of social information also depends strongly on the quality of information from the environment. This in turn has important consequences for the collective behaviour of social groups in different contexts. Different species of animal, for example, are habituated to widely differing levels of environmental noise. The same outcomes may have quite different fitness consequences for different individuals within an animal group. In human society too one finds contexts of varying uncertainty and agreement over preferences. The model predicts that these differing environmental and social contexts will lead to dramatically different social behaviour. Previous work has shown that rational individuals with identical preferences should exhibit similar social responsiveness across a wide range of environmental conditions [3]; in a noisy environment private and social information both become less reliable to the same degree, such that the balance between the two remains constant. Here I have shown that when preferences are not identical this symmetry is broken. In a low-noise environment, private information is more trustworthy than social information, since it is not corrupted by differences in utility function. In a noisy environment the uncertainty an individual has about others’ preferences is relatively low compared to the general uncertainty of all information, and thus the relative strength of social information increases. On the group level this should lead to greater consensus in collective decision making when individual preferences are strongly aligned and when uncertainty is high. These results imply that different species of animal (or the same species habituated to different environments) will display different social behaviour in the wild in a predictable fashion. Groups composed of individuals habituated to noisier environments should be more cohesive and responsive to the actions of others. In addition to a stronger overall social response, I also found that high noise environments and strong commonality in preferences led to a strong order-dependent use of social information, wherein a focal decision maker was more likely to follow a minority of recent decisions rather than a majority of preceding ones. This suggests that recent information will be dominant in cases where uncertainty about the world is generally high, and where a focal decision maker can assume that others want the same things as it does.

As noted in ref. [3], there is comparative ecological evidence to support the greater social responsiveness of agents habituated to noisier environments. However, this is complicated by the effect of context-dependency where animals have been tested under conditions that differ from their habitual environment, and the effect is likely to be amplified by studies in the laboratory under common conditions. In common with earlier predictions by ref. [3], here I found that reducing the experimental noise level relative to the habitual environmental level strengthened social response, increased consensus and made recent social information more important. Thus, individuals transposed from a noisy environment to the laboratory are subject to both the increased sociality imposed by adaptation to a noisy environment and also the heightened social responsiveness from a reduced experimental noise level. There is also likely to be a strong selection bias in the experimental literature within the field of collective behaviour towards instances where individuals display strong social interactions. Examples of trivially non-social behaviour in low noise environments, such as selecting the phone number for a known contact in a telephone directory, are therefore unlikely to be considered for study. These likely issues with existing experimental and observational data mean that further comparative studies are needed to assess how far the predictions in this paper are reflected in real populations.

Where agents are able to adapt their behaviour to a variety of different contexts, one can expect to see indi viduals change their social responsiveness as the situation demands. A consequence of this would likely be to induce strong social feedback and a high salience for the most recent social information when environmental uncertainty is high and individuals transparently share the same objectives. This is a possible contributing factor in several examples of destructive herding behaviour: stock markets consisting of agents with strongly aligned goals (investment returns) tend to eliminate the value of private information for most participants [32], leading to high uncertainty for the typical investor, conducive to bubbles and crashes; crowd disasters are often associated with situations where individual utilities are strongly aligned (to escape), as private information becomes less reliable (e.g. through the presence of smoke [33], or through alcohol consumption [34]). While in these instances such social feedback is ultimately maladaptive, such behaviour may be the result a social heuristic with a rational basis: when private information is scarce, follow others, especially those similar to yourself.

The results also demonstrate a potential source of leadership in heterogeneous groups. Where a group is composed of cryptic sub-types with differing preferences, we should expect those in the minority group to attend less to social information, and those in the majority to be more strongly social. On the individual level, this may contribute to the consistent differences in social response observed between individuals in some animal groups (e.g. [35, 36]). At the group level, it would create an emergent leadership role for those in the minority, who would attend relatively more strongly to environmental cues, while those in the majority act preferentially as followers and thus ensure group cohesion. Since cohesive groups can be led by a relatively small number of relatively less social individuals [20, 23, 25, 37], these minority groups could have a disproportionately large effect on the collective decision making. However, rather than the group being led by a subset of ‘informed individuals’, in this model leadership would be conferred on those with unusual preferences rather than greater information. Heterogeneity of preferences could result from genetic or ontogenetic causes, and one may speculate on whether these may be connected with ‘personality’ features such as boldness that have been linked to leadership [9, 38, 39]. Conflicting preferences could also result from transient circumstances such as different hunger levels, creating ‘leadership by need’ [25]. Phenotypic and behavioural heterogeneity plays a important role in how social groups function [40], and the potential role of minority-preference groups as leaders adds another mechanism whereby individual-level variation can be translated into group-level behavioural differences.

I also found that heterogeneous groups created a special role for unanimity in decision-making, especially in relatively low-noise environments. When private information is of high quality (low noise), individuals would ordinarily tend to follow this information rather than attend to the choices of others, unless preference alignment is extremely high. However, in groups where some individuals have near-identical preferences, the existence of a consensus among previous decision-makers can convince the focal individual to follow this consensus quite strongly (thus also reinforcing the consensus). Not only does this provide a mechanism for generating surprising degrees of consensus in groups that may have a low degree of average preference alignment, it also makes any deviation from that consensus especially powerful. Such a deviation from previous unanimity ‘breaks the spell’ and drives further decision-makers back to trusting their own private information.

It is important to reiterate that the model developed in this paper considers only the information value of other agents decisions. Aggregation may also carry intrinsic benefits [41], such a predation dilution [4], mate availability and temperature regulation [42]. These additional factors complicate the identification of social information use in real animals, especially in the wild [43]. Nonetheless, the results here suggest concrete predictions regarding collective behaviour: (i) stronger social interactions between individuals with similar preferences; (ii) greater aggregation in noisy environments; (iii) stronger salience of recent social information in the presence of uncertainty; and (iv) a correlation between leadership and minority types in heterogeneous groups. These predictions are potentially testable through both through comparative analysis in wild populations or through intervention studies in the laboratory, especially using artificial conspecifics [44–46] (though bearing in mind the prediction that social responses may be systematically stronger under laboratory conditions [3]). The effect of genuinely differing preferences (as opposed to differing information) on collective decision making has received relatively little experimental attention. The results in this paper show how future work may test whether any such effects are based on rational individual decision-making principles.

## METHODS

### Utility functions

I assume that any two possible choices can be distinguished by a set of values, *x*_1_,*x*_2_,…*x_n_*, characterising the difference between the options across a set of independent factors. I further assume that the utility difference, *U_k_* between the two options for any individual *k* is a weighted sum over these values:

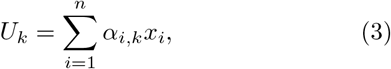

where the *α_i,k_* are the weighting coefficients for individual *k* and thus specify their utility function.

I assume that individuals are familiar with making decisions within a habitual environment and thus with the general properties of how the factors *x*_1_,…, *x_n_* vary between decisions. I further assume without loss of generality that each factor *x_i_* is measured on a standard scale, with mean zero and unit variance. Therefore, if the number of factors is large, or if the factors are themselves normally distributed, the prior distribution over the utility difference for individual *k* follows a normal distribution:

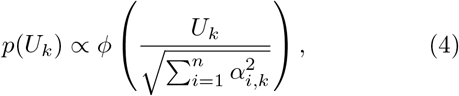

where *ϕ*(·) is the standard normal probability density function. Although individuals may have differing utility functions, I assume that these operate on the same scale, which I set without loss of generality to one. This is equivalent to specifying that 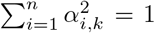 for all *k*, and therefore I retrieve the same prior distribution over utility differences as specified in [3]:

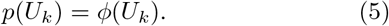

Through this definition of the utility function, one can also specify the joint distribution of utilities for multiple agents as a multivariate normal distribution 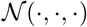:

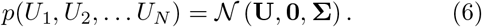

where **U** = [*U*_1_,…, *U_N_*]^T^, and

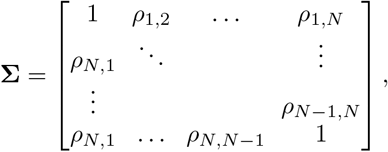

with *ρ_k,l_* being the correlation between any two utility functions: 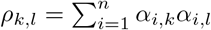. At this point I introduce the concept of an undifferentiated population. In such a population I assume that all pairs of agents are equally aligned in their preferences, averaged over possible choice characteristics *x_i_*, which corresponds to setting *ρ_k,l_* = *ρ* for all pairs *k, l*. This corresponds to a social environment in which agents know that others have somewhat differing preferences, and know the general degree of overlap between those preferences and their own, but do not keep track of which individuals may be more or less closely aligned to themselves.

### Private information

Each agent receives private information about the values of *x_i_* from the environment, for example through physical sensory mechanisms. This information is assumed to be imperfect, and corrupted by noise residuals *ν_i,k_* which are independent between individuals and between factors, such that the measured value of *x_i_*, denoted as 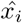, takes the form:

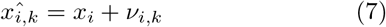

For convenience, I define a ‘privately estimated utility’ for agent *k, Û_k_*:

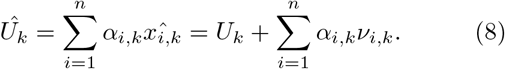

Defining *ϵ* as the environmental noise level relative to the scale of utilities, with 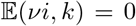 and *var*(*ν_i,k_*) = *ν*^2^, one can therefore specify the conditional probability distribution of the privately estimated utility for agent *k*:

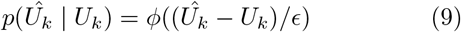

Taken alongside equation 6, this implies that the true and privately estimated utilities for all agents are jointly normally distributed, with:

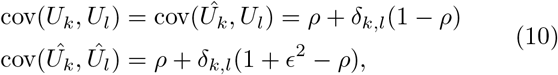

where *δ_k,l_* is the Kronecker delta function.

In the case of the first individual to make a decision, their belief about the relative utilities of the two options is entirely determined by the combination of their prior expectations and their private information:

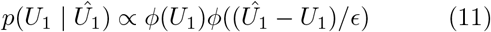

This implies that the first individual will choose option A if and only if (iff) *Û*_1_ > 0.

### Social information

Having observed the decision made by the first individual, *C*_1_, the second decision maker must combine this social information with their private information (*Û*_2_) to update their belief over the relative utilities based on their own preferences, *U*_2_. Recalling that the true and privately estimated utilities of all agents are jointly normally distributed, the second individual’s beliefs are updated as:

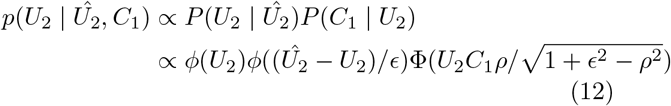

This belief over *U*_2_ further implies a critical value of the privately estimated utility, *Û*_2_* such that:

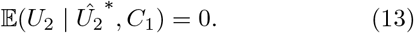

If *Û*_2_ > *Û*_2_^*^ then the second individual will choose option A, otherwise they will choose option B.

### Further decisions

The third decision maker can observe the social information from the first decision available to the second agent. Since I assume that agents are undifferentiated and share a common pairwise preference alignment *ρ*, the third decision maker can also determine the critical value *Û*_2_^*^ that is calculated by the second individual as above. Having observed the second decision, they are therefore able to determine whether the privately estimated utility of individual 2 is greater or less than this critical value. The third individual should then combine their own private information, summarised by their privately estimated utility *Û*_3_, with the knowledge they have of the bounds on what individuals one and two have observed. For example, if the third individual observes the first decision in favour of option A and the second in favour of option B, they should update their belief as follows:

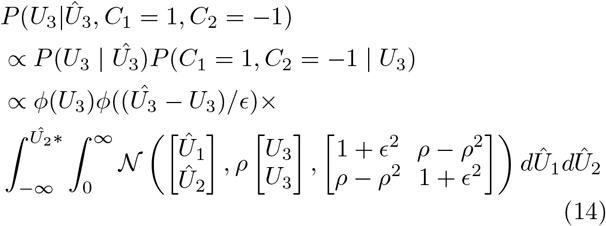

This updated belief structure further specifies a critical value 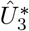 such that 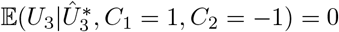. Subsequent agents can thus determine a bound on *Û*_3_ from the choice of individual 3, and this process can be followed recursively in turn for each further decision maker, with the limits of integration chosen based on whether each individual chose option A or B.

### Populations with cryptic sub-types

So far I have considered groups of undifferentiated individuals, making the approximation that every pair of individuals has the same degree of preference alignment. Now I consider groups drawn from a population in which there are two cryptic sub-types: individuals have a strong preference alignment with those of the same sub-type but a weak preference alignment with those of the other type.

That these sub-types are cryptic implies that the focal decision maker is not aware of the specific type identities of previous decision makers, only the overall prevalence of the two types in the population.

Consider a population composed of two types, *α* and *β*, and let *γ* be the proportion of the population that is of type *α*. I assume that individuals of the same type have a preference alignment of *ρ_α,α_* = *ρ_β,β_* = *ρ*_high_, while those of different types have *ρ_α,β_* = *ρ*_iow_, with *ρ*_iow_ < ρ_high_. If the focal decision maker can determine the types of the previous decision makers, then it can apply the model previously described, but with an adapted covariance between different individuals:

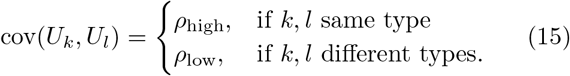

To determine its own utility belief function, a focal decision maker must evaluate the probability that the observed decisions were made, based on each possible sequence, *s* of types among previous decision makers, and weight these by their relative probabilities:

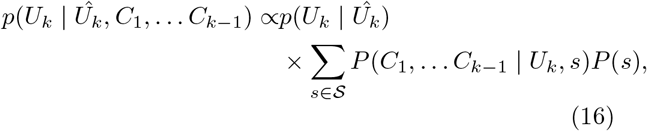

where 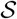 represents all possible sequences of individual types, and *P*(*s*) = *γ^nα^* (1 – *γ*)*^n_β_^*, with *n_α_* and *n_β_* being the number of individuals of each type in the sequence s. This process can be carried out recursively as for the undifferentiated population, with the focal agent calculating a different critical value of 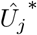 for a previous decision maker *j* depending on whether that agent was type *α* or *β* in a given putative sequence. Since the types of each individual are cryptic, the focal agent knows that other individuals are also ignorant of the types of the earlier decision makers.

## Acknowledgements

Graham Budd provided valuable feedback on the manuscript. RPM is supported by a UKRI Future Leaders Fellowship MR/S032525/1

